# Modelling the effects of adult emergence on the surveillance and age distribution of medically important mosquitoes

**DOI:** 10.1101/2025.04.10.648138

**Authors:** Isaac J. Stopard, Ellie Sherrard-Smith, Hilary Ranson, Kobié Hyacinthe Toe, Jackie Cook, Joseph Biggs, Ben Lambert, Thomas S. Churcher

## Abstract

Entomological surveillance is an important component of mosquito-borne disease control. Mosquito abundance, infection prevalence and the entomological inoculation rate are the most widely reported entomological metrics, although these data are notoriously noisy and difficult to interpret. For many infections, only older mosquitoes are infectious, which is why, in part, vector control tools that reduce mosquito life expectancy have been so successful. The age structure of wild mosquitoes has been proposed as a metric to assess the effectiveness of interventions that kill adult mosquitoes, and age grading tools are becoming increasingly advanced. Mosquito populations show seasonal dynamics with temporal fluctuations. How seasonal changes in adult mosquito emergence and vector control could affect the mosquito age distribution or other important metrics is unclear. We develop stochastic mathematical models of mosquito population dynamics to show how variability in mosquito emergence causes substantial heterogeneity in the mosquito age distribution, with low frequency, positively autocorrelated changes in emergence being the most important driver of this variability. Fitting a population model to mosquito abundance data collected in experimental hut trials indicates these dynamics are likely to exist in natural *Anopheles gambiae* populations. Incorporating age structuring into an established compartmental model of mosquito dynamics and vector control, indicates that the use of mosquito age as a metric to assess the efficacy of vector-control tools will require an understanding of underlying variability in mosquito ages, with estimates affected by short-term and seasonal fluctuations in mosquito emergence.

## 1 Introduction

Entomological surveillance is an important component of mosquito-borne disease control and is regularly used to assess entomological risk and the effectiveness of current and potential future vector control interventions [1, 2]. Mosquito abundance, parity (the proportion of mosquitoes that have laid eggs), the proportion of mosquitoes that are infectious and the entomological inoculation rate (rate a which people receive infectious bites) are currently the most widely used metrics. These metrics are, however, difficult to measure accurately [1], with mosquito catches depending on the location and trapping methods [3], and noise in adult mosquito emergence making it difficult to infer changes in transmission [4]. Indeed, epidemiological and entomological end points of vector control trials do not always indicate effects in the same direction [2].

To transmit malaria-causing *Plasmodium* parasites, and numerous arboviruses, mosquitoes must survive the extrinsic incubation period (EIP). For many mosquito-borne diseases the EIP is long relative to mosquito life expectancy [5, 6], meaning small changes in the adult mosquito age distribution cause large changes in entomological transmission intensity [7–9]. This explains, in part, why interventions that reduce mosquito survival have been so important to mosquito-borne disease control [7, 10, 11]. Measuring the age of wild mosquitoes is challenging [12, 13]. Historically, the age distributions of *Anopheles* mosquitoes were estimated using morphological methods: the physiological age, for example, was estimated by dissecting mosquitoes and counting the number of dilations at the base of the ovarioles to estimate the number of gonotrophic cycles that the mosquito has survived (Polovodova’s technique) [13]. Calendar age can then be approximated by making an assumption about the length of the gonotrophic cycle, which is generally assumed to be about 3 days [14]. Alternatively, calendar age can be measured morphologically by counting the layers of cuticular growth on the inner apodemes, which increase daily [15]. These methods are, however, laborious and require substantial skill, meaning they are impractical to use on a large scale [12]. Novel age grading techniques that measure calendar age using spectroscopic or gene and protein profiling methods are in development [12], and studies employing spectroscopy and machine learning models have recently demonstrated increases in accuracy [16, 17]. Wild mosquito age distribution data may therefore become more available in the future.

Age grading of wild mosquitoes has been proposed as a method to evaluate novel vector control tools which kill adult mosquitoes: as increased mortality could be indicated by a shift in the mosquito age distribution [12]. Although intuitive, this has not been directly observed, either through mechanistic simulation models or in field data, though changes in mosquito parity (a proxy for age) have sometimes been observed [18]. Hypothetically, the age structure of the mosquito population will be driven by changes in both adult mosquito mortality and emergence; during periods of increased emergence, for example, the proportion of young mosquitoes increases, which means the temporal dynamics of mosquito abundance can affect the mosquito age distribution [19]. These processes therefore also affect parity and infection prevalence, which depend on mosquitoes surviving the first gonotrophic cycle or EIP respectively.

Medically important mosquito populations often show seasonal dynamics and spatiotemporal fluctuations [20, 21], contributing to seasonal patterns of transmission [22]. Population dynamics can be driven by demographic and environmental processes: vital demographic rates, for example, can change due to changes in environmental variables or intrinsic demographic processes (such as density dependence) [23–25]. Changes in demographic rates can occur at fixed frequencies resulting in seasonal population dynamics (seasonal or periodic forcing). These temporal trends in the population are classed as red when low-frequency, positively autocorrelated changes dominate (for example, high emergence today is likely to be followed by high emergence tomorrow), blue when high-frequency, negatively autocorrelated changes dominate (high emergence today is likely to be followed low emergence tomorrow) and white when no frequency dominates [26]. Empirical temporal abundance data spanning more than 30 years for 123 species indicates that changes in abundance are typically low-frequency, though smaller animals tend to have less red fluctuations [27]. The red frequency of temporal fluctuations in the abundance is commonly attributed to changes in environmental variables, because positive autocorrelation is observed in both the abundance and environmental variable time-series [28]. Species with high growth rates theoretically respond more to the colour of environmental noise, meaning they are more sensitive to differences in the autocorrelation of environmental variables [26, 29]. Indeed, anopheline mosquito dynamics show species-specific responses to certain environmental variables, such as rainfall, temperature, humidity, windspeed and land use [21, 22, 30–34]. Fluctuating mosquito populations in time and space will likely increase the noise in entomological metrics and could explain some of the high variability observed in field data, complicating interpretation [2]. There has, however, been little research on the colour of temporal fluctuations in adult mosquito emergence and the resultant impact on the variability of adult mosquito metrics and age distribution.

In this study we aim to (1) develop a mathematical model of adult mosquito population dynamics that explicitly allows quantification of the mosquito age distribution and parameterise it for the dominant malaria vector *An. gambiae*, (2) investigate how temporal heterogeneity in the age distribution of the mosquito population is affected by differences in the frequency and autocorrelation of fluctuations in mosquito emergence, (3) estimate the observed autocorrelation in the emergence of wild *An. gambiae sensu lato* mosquitoes and predict changes in the age distribution given these changes in emergence, and (4) investigate the impact of vector control on the mosquito age distribution with variable emergence.

## 2 Results

### 2.1 Mosquito emergence can drive patterns in mosquito metrics

We developed a stochastic individual-based model (IBM) of female adult *An. gambiae* mosquito dynamics that tracks individual mosquito emergence, parity, aging and mortality over discrete time steps. In this model, mosquito survival, emergence and gonotrophic cycle duration are treated as random variables represented by probability distributions, and the model is parameterised assuming a constant per-capita mosquito mortality rate. In this model adult mosquito emergence is stochastic; the numbers of adult mosquitoes emerging varies daily following a Poisson distribution (o_*t*_ *∼* Poisson(*λ*_*t*_)) and the mean emergence rate (*λ*_*t*_) can vary temporally, with the variation characterised by either (a) different frequencies (represents the outcome of periodic forcing, captures the number of transmission seasons per year) or (b) different levels of temporal autocorrelation (correlation in emergence between successive days). This model is used to estimate both temporal changes in mosquito abundance, parity and the age distribution of a group of adult mosquitoes over time (Figure 1A). The parity and age distribution of mosquitoes is assessed from all simulated mosquitoes with approximately 200 mosquitoes initially and we calculate the metrics using all simulated mosquitoes.

**Fig 1.**
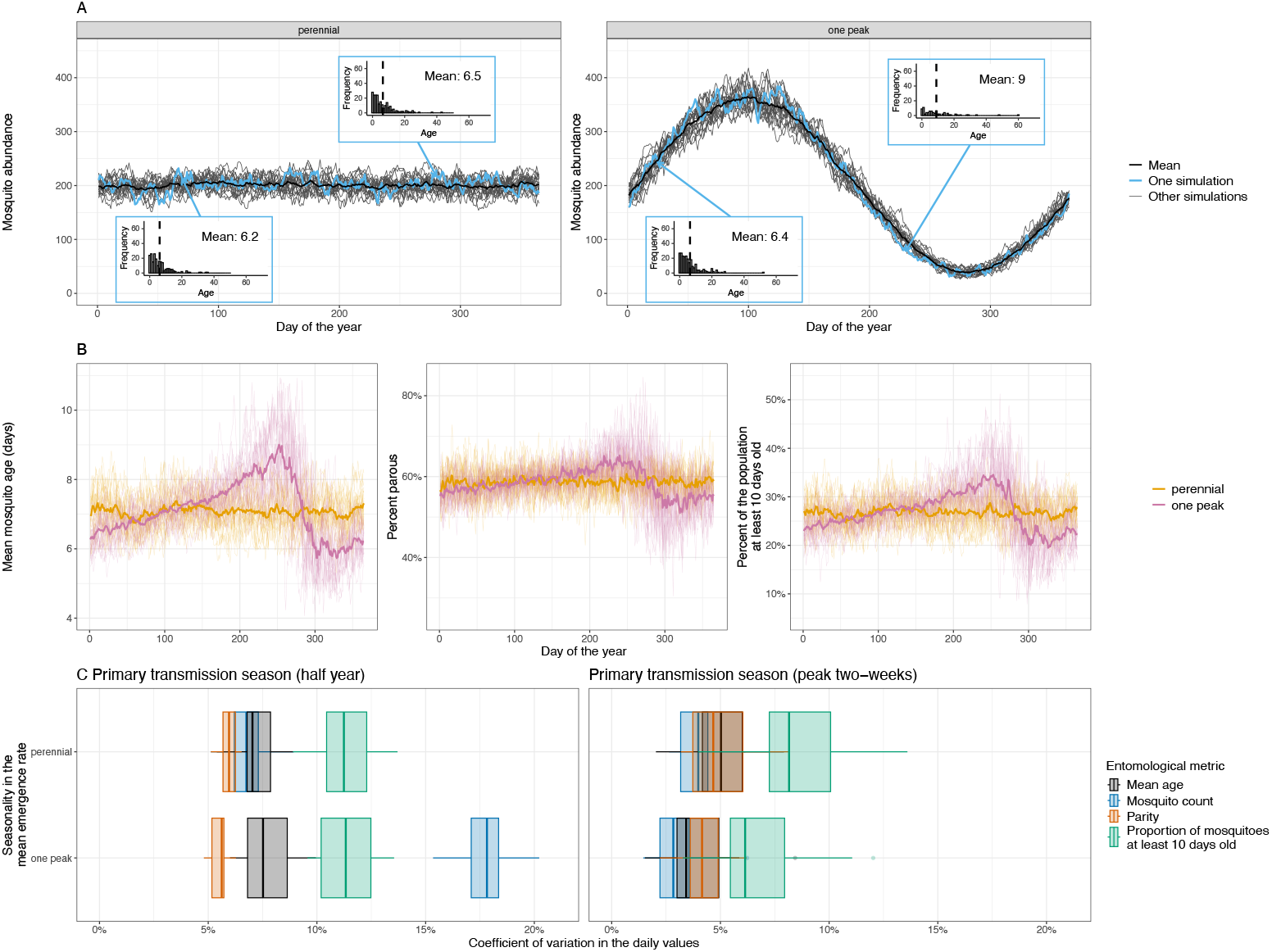
Seasonal dynamics in mosquito emergence changes the adult mosquito age distribution. (A) The temporal dynamics in mosquito abundance from single simulations assuming the mean adult, female mosquito emergence rate is either time-invariant (perennial dynamics) or follows a sine wave with a single period per year (one peak dynamics). For one of these simulations (highlighted in blue) we presented the predicted mosquito age distribution on two different days; in these histograms the dashed vertical line indicates the mean age. For the seasonal setting the day selected was when the mosquitoes abundance was increasing or decreasing. In the main plots the thicker black line show the means abundance value from all the simulations. The initial number of mosquitoes in the group was approximately 200. For all plots, the model was simulated with a an overall mean emergence rate of 26.4 per day over the year (equal to the mean number of mosquitoes that die per day). For the seasonal simulations the mean emergence rate was assumed to vary at a frequency of 1 rad per year with an amplitude of 20 mosquitoes. (B) shows how the mean mosquito age, percent of mosquitoes that are parous or at least 10 days old varied with time given different seasonal patterns in mosquito emergence. The thin lines show the results from a single simulation and the thicker lines show the mean values of all the simulations. (C) Shows how the coefficient of variation (ratio of the standard deviation to the mean) over the primary transmission season (day 1 to 180) and two-week centred on the peak transmission season varies depending on the entomological metric used. Box plots show the variability between different simulations, with the central values being the median, hinges showing the 25^th^ and 75^th^ percentiles, and the whiskers indicating the smallest or largest values no further than 1.5 interquartile ranges from the hinges.

Stochastic processes cause noise in the mosquito age distribution, but given perennial dynamics (a constant mean adult mosquito emergence rate) the modelled age distribution and mean age of the mosquitoes are observed to be relatively stable over time (Figure 1A & B). In contrast, when mean adult mosquito emergence varies seasonally the model produces seasonal variability in the mosquito age distribution, with it shifting towards younger mosquitoes when the abundance is increasing, and older mosquitoes when the abundance declines (Figure 1A & B). The range in the mean daily mosquito age over a single year is substantial; for example, it varied over the year between 4.1 and 14.0 days in simulations with a one peak transmission season and 5.3 and 8.8 days in those with perennial dynamics (Figure 1B). Similar patterns are seen in the proportion of mosquitoes that are parous (Figure 1B). Greater variability is predicted in the proportion of mosquitoes at least 10 days old, analogous to the proportion of mosquitoes capable of being infectious with malaria, fluctuating between 8% to 51% in all simulations with seasonal emergence, and 17% and 39% in those with perennial dynamics (Figure 1B). We calculatec the coefficient of variation (ratio of the standard deviation to the mean, CV) of the temporal changes if the different entomological metrics (Figure 1C). In simulations with perennial dynamics the CV of both mosquito abundance and the mean mosquito age are similar irrespective of the time frame considered (either the complete primary transmission season [half year] or the two weeks when abundance peaks) (Figure 1C). When the time-frame considered includes the complete primary transmission season (180 days) seasonality in mosquito emergence increases the CV in the mosquito abundance, but not the other metrics (Figure 1C). When the time frame considered is shorter (the two-weeks that includes the peak of transmission) the CV in mosquito abundance is less and the CV for the proportion of mosquitoes older than 10 days is the most variable metric (Figure 1C). To assess the robustness of these results, we investigated the sensitivity of temporal changes in the mean mosquito age to differences in the (1) frequency (assuming a medium amplitude) (Figure S1) and (2) amplitude (assuming one peak per year) of fluctuations in the mean adult mosquito emergence rate (Figure S2). The CV in the mean daily mosquito ages was most sensitive to larger amplitude changes in adult mosquito emergence at low to medium frequencies (approximately 5-10 radians per year) (Figure S3A & B).

Age grading has been proposed as a method of estimating the mosquito mortality rate. We investigated how variable emergence might bias these estimates by fitting a survival model to the simulated mosquito age distributions from each day. Individual mosquito survival is treated as a Bernoulli trial in the model, with the probability a mosquito dies assumed to be constant with time. This model assumes that the mosquito population is stable and age distribution mirrors the life time distribution. This is not the case in the simulations with seasonal emergence, meaning the fitted per-capita mortality rates were biased, the values from all simulations, for example, varied between 0.11 and 0.19 over the year given perennial changes in mosquito emergence and 0.07 and 0.25 given a one peak transmission season (Figure S4). These methods can therefore not accurately estimate mortality in seasonal settings.

### 2.2 Estimated changes in the emergence of wild mosquitoes

Temporal changes in the adult mosquito emergence rate can be characterised by the level of autocorrelation. Theoretically, the lag-1 autocorrelation can be positive (*h* > 0), meaning there is positive correlation between the rate adults emerge on subsequent days, neutral (*h* = 0), or negative (*h* < 0), indicating that if the abundance of emerging adults is high one day it is likely to be low the next (Figure S5A). This range of temporal autocorrelation is explored with the model to see how the autocorrelation influences variability in the mosquito abundance (Figure S5B) and age distribution (Figure S5C). Hypothetically, positive autocorrelation (red fluctuations, high seasonality) increase the variability in daily estimates of the mean mosquito age over a single year more than negatively autocorrelated (blue) fluctuations (Figure S5C, Figure S6, Figure S7).

To estimate changes in the emergence of wild mosquitoes, we fitted a simplified ordinary differential equation (ODE) version of the adult mosquito model, which models changes in mosquito abundance assuming a time-varying daily mosquito emergence rate (characterised by a lag-1 autoregressive model) and constant mortality rate, to previously published, temporally disaggregated experimental hut trial (EHT) data from Burkina Faso [35] (Figures 2A & B). The ODE model posterior mean emergence rate, lag-1 autoregressive model standard deviation, and overdispersion parameter values are shown in Figure S8. The estimated posterior autocorrelation parameters in the daily emergence rate exceed zero, indicating that mosquito emergence is positively autocorrelated (Figure 2C).

**Fig 2.**
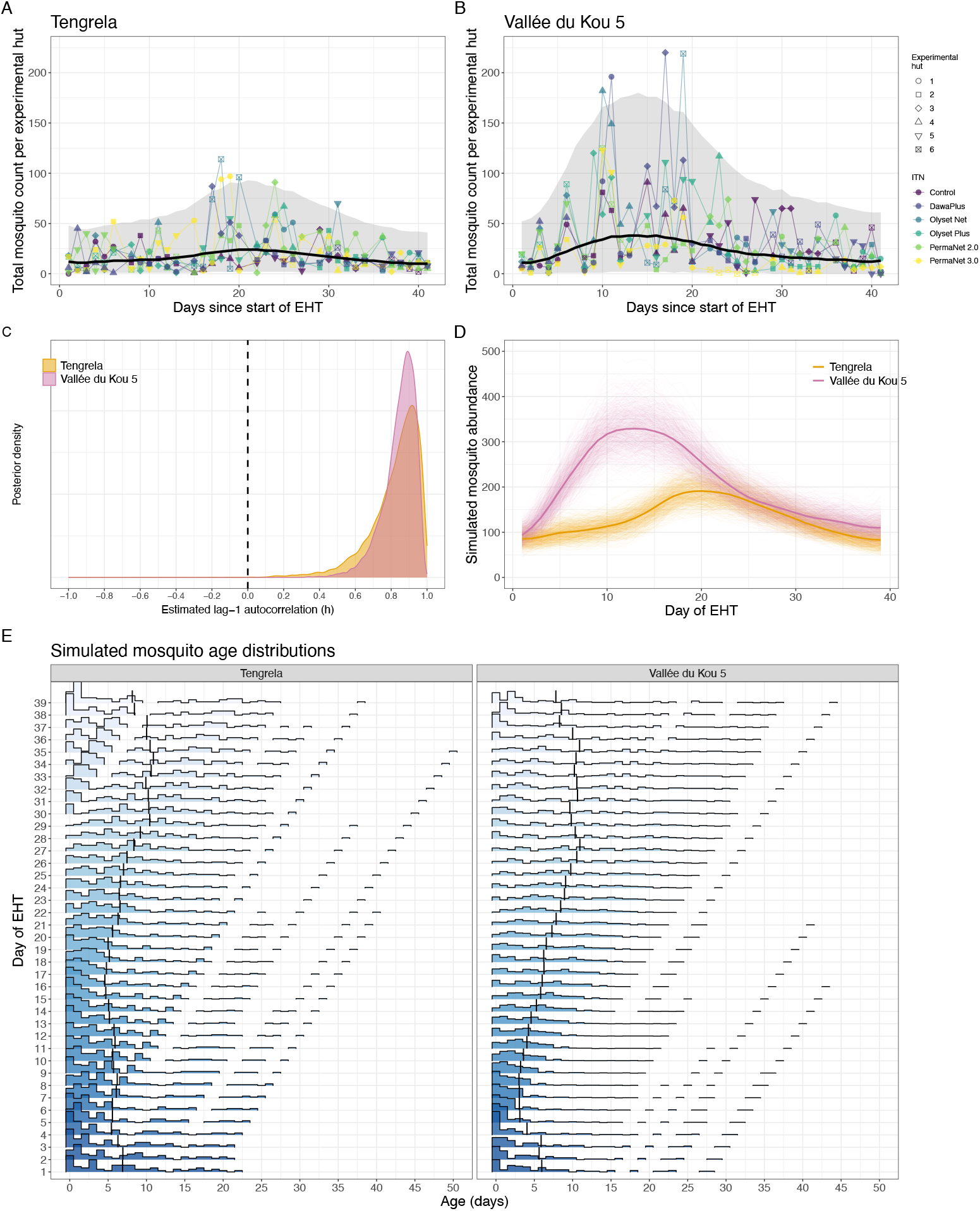
Changes in the adult mosquito age distribution predicted in experimental hut trials (EHTs). (A) and (B) the numbers of mosquitos captured during the EHTs. The line shows the number of mosquitoes predicted by the ordinary differential equation (ODE) model. The ODE model is fit to all experimental huts and gives the number for a single experimental hut. (C) The estimated autocorrelation in the adult mosquito emergence rate (λ*t*) in the ODE, assuming a constant per-capita mortality rate. (D) The simulated mosquito abundance in the individual based model using the fitted *λt* values as the mean of the Poisson distributed numbers of adult mosquitoes emerging. The initial numbers of mosquitoes (*M*_0_) and *λ*_*t*_ values were multiplied by the number of experimental huts to give the total abundance across all experimental huts. The thin lines show the values for 750 simulations given different samples from the posterior parameter values. The thick lines shows the mean simulated abundance. (E) The predicted changes in the mosquito age distributions during the EHT for a single random simulation with one random draw from the posterior *λt* estimates. EHT data was obtained from [35].

To estimate potential changes in the age distribution, the estimated initial numbers of mosquitoes and daily adult mosquito emergence rates were multiplied by the number of experimental huts and used as parameter inputs within our IBM. The resultant simulated changes in mosquito abundance and the age distributions of the mosquito population during the EHTs are shown in Figures 2D and E respectively. Despite the short duration of the EHTs there is clear predicted temporal variability in the mosquito age distributions, with the predicted mean mosquito age varying between 4.6 and 10.9 days in the Tengrela EHT and 3.0 days and 10.9 days in the Vallée du Kou 5 (VK5) EHT, with the mosquitoes being younger during the start of the EHT and older towards the end (Figure 2E).

### 2.3 Investigating the impact of interventions on mosquito age

To assess the effects of insecticide treated bed nets (ITNs) on the age distribution of the mosquito population, we developed a discrete time stochastic version of a well established compartmental, differential equation model of seasonal larval and adult mosquito population dynamics that includes vector control interventions [36, 37]. We adapted this model to track mosquito age by transitioning mosquitoes between different age compartments with each discrete time step (*dt* = 1 day). We simulated seasonality in mosquito population dynamics (implemented through changes in the larval carrying capacity) based on either (1) rainfall for the Cascades region of Burkina Faso or (2) perennial dynamics (Figure 3A). Pyrethroid-only ITNs were introduced when the mosquito population was either increasing or decreasing. The model indicates ITNs consistently decrease the mean mosquito age, irrespective of the implementation timing (Figure 3B). For example, in a perennial setting, ITNs are predicted to reduce mosquito abundance by 63% (mean value from all simulations, ranges from 50% to 75%) after the first month and reduce the mean age from 7.2 days (mean value from all simulations, ranges from 6.1 to 8.4 days in all simulations) to 2.2 days (ranging from 1.6 to 3.4 days, a drop of 69% [range in all simulations: 55-80%]). Similar patterns were seen if the intervention was implemented early in the transmission season, reducing mosquito abundance by 62.2% (range from 50.4% to 71.4% in all simulations) and reduce the average age from 6.4 days (5.8-7.3 days in all simulations) to 2 days (1.4-2.7 days in all simulations, a drop of 68.7% [58.3-76.7% in all simulations]). However, implementing the same intervention towards the end of the transmission season the model predicts a 63.8% (ranging from 52.3 to 74.7% in all simulations) reduction in mosquito abundance after the first month, with the average age reducing from 11.4 days (ranging from 8.2 to 15.3 days in all simulations) to 2.9 days (1.5 - 4.3 days in all simulations, a drop of 74.5% [62.1-88.2% in all simulations]). When the mosquito abundance is decreasing the mean age increases and once ITNs are implemented the magnitude of these fluctuations decreases (Figure 3B). This means that if ITNs are rolled out when the mosquito population is decreasing, the effect of the ITNs on the mean adult mosquito age may appear larger than if ITNs (or other vector control) are rolled out as vector abundances increases (Figure 3C). This demonstrates the importance of understanding age structure dynamics to interpret intervention effects and the importance of when sampling is conducted.

**Fig 3.**
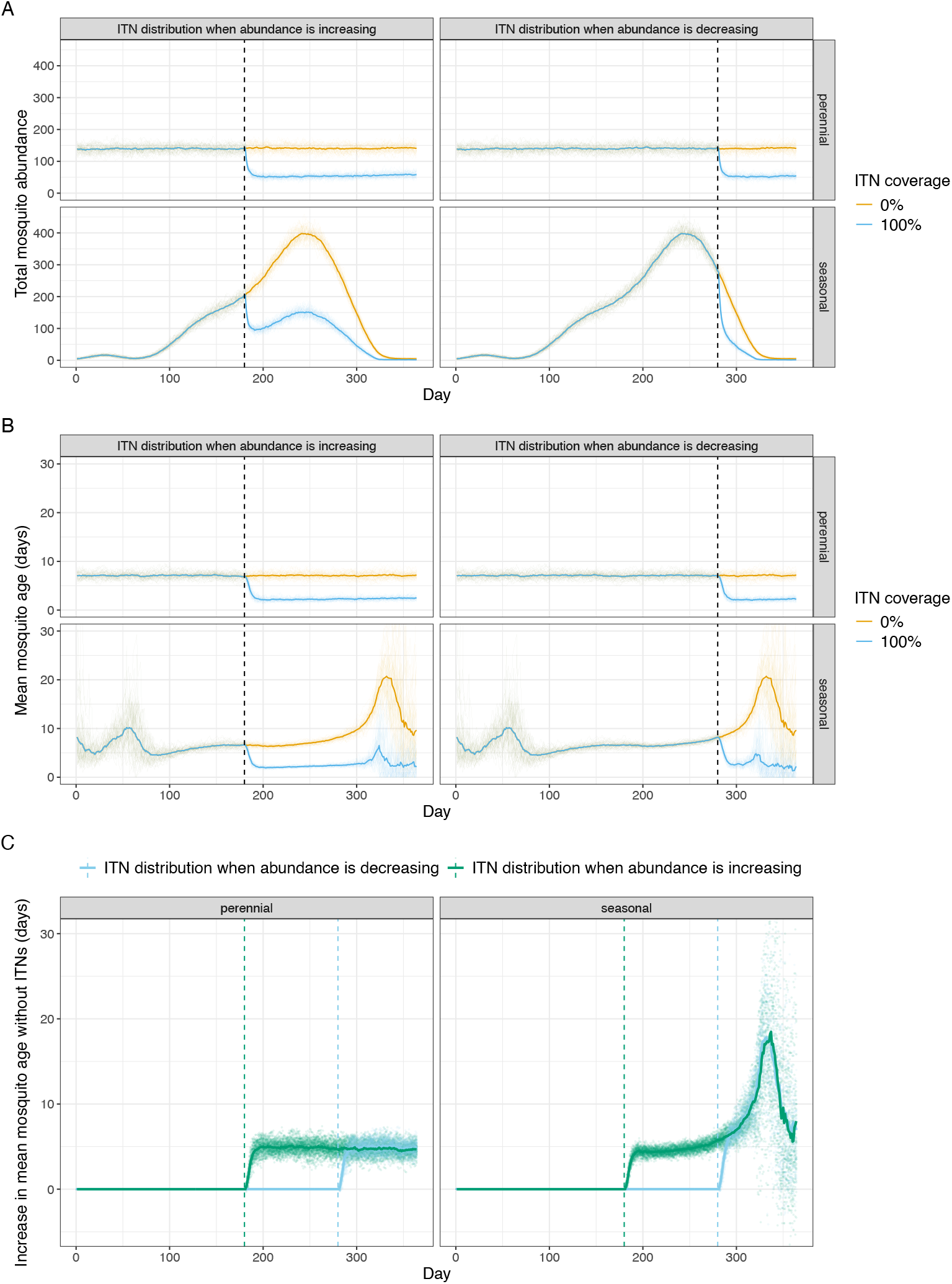
The effects of insecticide treated nets (ITNs) on the age distribution of the mosquito population. Mosquito population dynamics are simulated using a state-space version of an established compartmental mosquito population model [36, 38], with seasonality determined by either (1) perennial dynamics or (2) rainfall in Cascades region of Burkina Faso. Panels (A) shows estimates of mosquito abundance, (B) the mean age of the mosquito population, and (C) the difference in the mean age of mosquito populations with or without ITNs. In A-B the yellow lines indicate simulations with no ITNs, with blue lines denoting populations where ITNs are deployed at the vertical dashed line (day 180 of the year for the panels titled “ITN distribution when abundance is increasing” and day 280 in panels titled “ITN distribution when abundance is decreasing”). Multiple stochastic realisations of the model are shown, with the mean value denoted by the thicker line.

## 3 Discussion

There has been substantial research into the theoretical importance of the mosquito age distribution for disease transmission [7, 8] and consequently also methods to estimate the age of a mosquito [12, 13, 16, 17, 39]. Here, we use mathematical models to demonstrate how the age of mosquitoes could change in response to variable emergence (noise and seasonal trends in abundance) and the use of control interventions. The model is parameterised using field data from the most important vector of malaria and demonstrates how conventional mosquito control is likely to cause substantial shifts in the age distribution of the mosquito population. Simulations like these can be used to support the development of mosquito age grading methods by demonstrating the level of precision needed for a technique to have practical use as surveillance tools. The work highlights that interpreting changes in the age distribution of wild mosquitoes is complex as there is likely to be considerable noise and seasonal trends in the age distribution of mosquitoes. Assuming all else is equal, the IBM of *An. gambiae* population dynamics predicts that annual seasonality (low frequency, highly autocorrelated changes in mosquito emergence), such as those identified in Burkina Faso, will likely cause the greatest temporal variation in mosquito ages. It has been suggested that age structures could be used to assess the impact of vector control interventions by evidencing different distributions in control and treatment settings during trials [12]. This modelling exercise suggests that we would need sufficient *a priori* information about seasonality to interpret results for this purpose. Indeed, in our compartmental model with vector control, if the changes in the age distribution are just measured pre- and post-intervention, the apparent change in the mosquito age distribution due to ITNs can be affected by when the ITNs are distributed relative to the transmission season. Interpreting changes in the mosquito age distribution will therefore need to carefully consider the timing of sample collection and be aware of seasonality and differences in the local fine-scale variability between experimental sites depending on, for example, proximity to breeding habitats [8], which could bias results. However, simulations suggest that the variability in the mean age of mosquitoes is likely to be less than that observed in entomological metrics currently used such as abundance and infectiousness (Figure 1C). This finding leads on from previous work that demonstrated the uncertainty around these conventional metrics caused by variable mosquito emergence [4]. This variability is likely to be impeding the statistical power of cluster randomised control trials with entomological endpoints. Models like the IBM presented here could be parameterised with field data to allow optimisation of sampling strategies and the generation of robust power calculations. For example, this work demonstrates considerable nightly variability in emergence in Burkina Faso, so mosquito collection over two nights apart may give a more reliable estimate of mosquito abundance, parity, sporozoite prevalence and mean age than samples collected over two consecutive nights. Further analysis of field data is needed to generate guidelines for improving the precision of entomological sampling.

We parameterised our model for *An. gambiae*, however, due to similar mechanisms these findings are likely to be relevant to other important vectors of mosquito-borne diseases. Sampled mosquito abundance for other mosquito species has been shown to vary at a range of frequencies: the Fourier transforms of daily light trap data for *Aedes vexans* and *Culiseta melanura* sampled over nine years indicate temporal changes in mosquitoes occur at frequencies corresponding to periodicities between two days and multiple years [40]. It has been suggested that generation times of mosquitoes mean that high frequency fluctuations in mosquito abundances of timescales of less than one month are, however, more likely to be due to behavioural responses [40], though care should be taken extrapolating between species due to differences in ovipostion behaviour and the type of breeding habitats.

The modelling exercise made a number of simplifying assumptions. First, it only considers heterogeneity in the emergence of adult mosquitoes and there are likely to be many other sources of variability that will impact mean entomological metrics and estimates and their associated uncertainty. For example, many mosquito demographic processes such as blood-feeding and mortality are likely to vary daily given local environmental conditions. This is likely to substantially increase the heterogeneity of different metrics outlined in Figure 1 and should be considered in sample size calculations. We do not consider biases in the trapping methodology, which could be substantial (particularly at young mosquito ages) and is likely to bias mean age, parity and infectiousness estimates. Second, in the IBM the way we model mosquito emergence means we assumed that increases in the sampled adult mosquito population due to aestivation and migration are equivalent to newly emerging adults (aged zero). Adult mosquito dispersal and the spatial distribution of larval habitats can, however, also influence population dynamics [41]. The age distribution of migrating mosquitoes is also, theoretically, important for mosquito infection dynamics: assuming a gradient in the mosquito age distribution with distance from larval habitat, for example, mean sporozoite prevalence would increase with dispersal distance [19]. When using the population-level emergence rate estimated from the EHTs to simulate changes in the mosquito age distribution we assumed all mosquitoes came from the same breeding site. Theoretically, the portfolio effect could, however, increase the stability of aggregate mosquito abundance due to the statistical averaging of stochastic fluctuations in the emergence of mosquitoes from different breeding sites [42]. Within the compartmental model we have accounted for potential feedback between the mosquito population size, age distribution and mosquito emergence, though many of the environmental and density-dependent processes governing mosquito emergence remain unclear so the reliability of the immature mosquito model needs further validation. In all models, we assumed a constant background per-capita age-independent mortality rate. Although age-dependent *Anopheles* mortality is observed in the laboratory [5, 43]; in the field where mosquitoes have much shorter lifespans senescence is likely to be less important [44]. Using Povolodova’s method, for example, Gillies & Wilkes [39] found a relatively constant mortality rate between mosquitoes with different physiological ages up to approximately the 7^th^ gonotrophic cycle. A meta-analysis of mark-release-recapture studies found little evidence of age-dependent mortality in multiple mosquito genera [45]. Mosquito mortality is, however, also strongly temperature and humidity dependent in the laboratory [5, 46], which we do not account for. There is empirical evidence for temporal variability in mosquito mortality in wild mosquitoes; Ngowo et al., for example, fitted a state-space model to temporal abundance data of multiple *An. funestus* life stages finding both the fitted survival and fecundity varied temporally [47]. Mark-release-recapture studies of male *An. gambiae*, however, found little effect of seasonality on mosquito mortality [48]. Differences in field and laboratory findings may exist due to mosquito resting behaviour and thermal avoidance [49–52], meaning the relationship between mosquito mortality and temperature, particularly the critical thermal limits, in the field remains an important question. Mosquito age distribution data from wild mosquitoes could therefore help identify the relative inputs of these different mechanisms, but our work demonstrates that interpreting changes in the age distribution of mosquito populations will require a mechanistic model that explicitly accounts for changes in both emergence and mortality. Briet [53], for example, developed methods for estimating adult mosquito mortality rates from parity data whilst accounting for differences in emergence. The model we develop could be fitted to a combination of age structure and abundance of different mosquito life stages to help (1) understand the importance of different mechanisms and (2) estimate sample sizes required to power studies investigating changes in mosquito mortality.

## 4 Methods

### 4.1 Data

EHT data were obtained from a single previously published study that carried out EHTs in both the villages of Tengrela and Vallée du Kou 5 (VK5) in the Cascades Region of Burkina Faso [35]. EHTs are an assay used to assess the entomological efficacy of vector control tools; volunteers rest within specially designed house-like structures with different interventions applied, all mosquitoes that enter the experimental hut are trapped and the numbers alive and dead are counted at the beginning of each day [54]. Data for mosquitoes identified as *An. gambiae s. l*. were analysed, though *An. coluzzi* was the only species identified by polymerase chain reaction (PCR) in a subset of these mosquitoes caught. Each EHT consisted of six experimental huts, with five insecticide treated nets (ITNs) and one untreated net tested. Study participants slept in the experimental huts six days a week. The study was conducted between the 8th of September and the 22 October 2014, with 36 sampling days (for one day per week there was no sampling). Differences between nets were ignored as the analysis used the number of mosquitoes caught per night and did not consider mosquito mortality or feeding status.

### 4.2 Individual based adult mosquito model

We developed the individual based model (IBM) to simulate adult female mosquito population dynamics over discrete time steps using the *individual* R package version 0.1.17 [55]. For all models and equations we assumed a time step of size *dt* =1 day, meaning the mosquito ages are limited to the nearest complete day. The model is stochastic and at each time step the number of emerging adult female mosquitoes, *o*_*t*_, is,

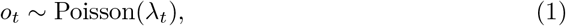

where *λ*_*t*_ > 0 varies over time *t* (in days). This model does, therefore, not allow feedback between the current mosquito population size and numbers of emerging adult female mosquitoes, but allows the impact of different types of variability in adult mosquito emergence to be isolated. Given *a* > 0 is the constant per-mosquito biting rate per day, then the probability a mosquito has bitten during a single day is given by the exponential distribution cumulative distribution function (CDF): 1 *e*^−*a*^. Newly emerged mosquitoes are all nulliparous (NP), and the number of simulation time steps for completion of the first gonotrophic cycle, *G*_*i*_, of each mosquito, *i*, is sampled,

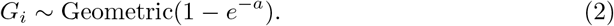

Once the first gonotrophic cycle is complete mosquitoes transition to the parous, *P*, state. Following existing *Anopheles* population models and empirical inference [45], mosquitoes are assumed to die at a constant age-independent mortality rate, *μ*, such that at each time step individual mosquito survival, *S*_*i*_ is determined by,

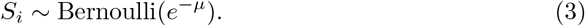

Dead mosquitoes are removed from the population, and at each time step the age of the individual mosquitoes is incremented by *dt* = 1. In existing continuous time malaria transmission models [37] it is assumed that the biting rate and mortality rate prior to the introduction of control interventions for *An. gambiae* are 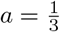 and *μ* = 0.132 respectively. In the continuous time model where biting times and adults mosquito life expectancies are exponentially distributed these give mean times of 3 and 7.6 days respectively. Using the geometric distribution and binomial approximation to transition mosquitoes between time steps gives mean biting times and life expectancies of 2.5 and 7.1 days.

#### 4.2.1 Simulation

The daily variability in adult mosquitoes emergence will depend on the geographical scale being considered in addition to the oviposition behaviour and bionomics of the mosquito (for example, whether a mosquito skip-oviposits or lays eggs in clumps). Variability in emergence from a single breeding site may be considerable, but this heterogeneity could be dampened due to statistical averaging if adult mosquitoes from multiple distinct breeding sites are caught in the same trap (the portfolio effect) [42], which is not considered in our analyses. The impact of stochastic fluctuations will also depend on the size of the mosquito group considered. Here we consider a theoretical, dynamic group of adult female mosquitoes. To represent the underlying population structure the model outputs the age distribution of living mosquitoes at all time points, which is equivalent to collecting and age-grading all mosquitoes. We do not consider heterogeneity generated by the sampling process, for example caused by trapping biases or small sample sizes.

To assess the hypothetical impact of intra-annual changes in mosquito emergence, we characterised temporal changes in the adult mosquito emergence rate, *λ*_*t*_, usinga sine function;

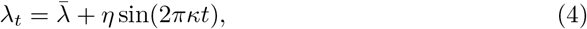

where 2*πκ* is the angular frequency, *ε* is the amplitude and 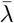 is the time invariant mean of the mean numbers of mosquitoes emerging. To ensure the population does not go extinct or increase exponentially, we set the mean number of births per day equal to the daily mortality probability multiplied by the initial number of mosquitoes (*M*_0_); 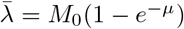. This ensures an average of 200 mosquitoes over each stimulation, though the number of mosquitoes changes over time according to the seasonality under consideration. The number of mosquitoes is approximately informed by the EHT data, where a median of 16 mosquitoes were caught per experimental hut during the transmission season, there are 6 experimental huts and it is assumed mosquitoes are only caught when attempting to blood-feed.

We assessed temporal changes in the mosquito age distribution and parity given the following seasonality profiles:

1. no long-term changes in emergence (*κ* = 0), which is representative of perennial mosquito abundance, and,
2. a single peak per year 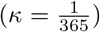, which is representative of wet-dry season dynamics.

To investigate the sensitivity of the modelled age distribution to different amplitude fluctuations in mosquito emergence, for the single peak per year seasonality profile we simulated mosquito emergence with a range of amplitudes (*η*). To assess the effect of different phenomenological changes in mosquito emergence on the temporal variability in the daily mean mosquito age, we ran the model with a range κ values corresponding to a frequencies between 1 and 50 radians per year with an amplitude (*η*) of 10. For these seasonal trends the sine wave frequencies corresponded to complete periods over a single year.

To model the effects of temporal variation in the mean adult mosquito emergence rate due to stochastic noise, for example due to hypothetical variation in a environmental variable [28], we applied a discrete-time lag-1 autoregressive model,

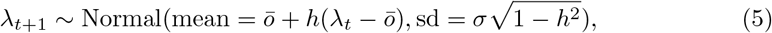

where, 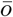 is the time-invariant fitted mean emergence rate (also assumed to be equal to *M*_0_(1 − *e*^−*μ*^)), *h* is the autocorrelation parameter and *σ* is the standard deviation. We generated stochastic changes in the emergence rate for the duration of the simulations for numerous values of −0.9 ≤*h* ≤0.9 with a 1-day lag and three values of *σ* ∈ (1, 3, 5). The 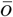 and *σ* values were selected such that *λ* was greater than zero.

For each parameter set for the frequency and autocorrelation sensitivity analyses the mosquito population dynamics were simulated 20 times for 5 years and the first 4 years excluded. For each simulation we calculated the temporal variability in the mean mosquito age, abundance, proportion of mosquitoes at least 10 days old and parity over the final year, and quantified temporal variability in these values using the 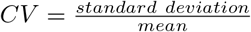.

To estimate the per-capita mortality rate from the mosquito age distributions, we added 10^*−*5^ to each age, such that no mosquitoes were aged zero, and fit a parametric survival model to the mosquito ages using the R survival package [56], assuming a constant per-capita mortality rate, exponential distribution and mosquitoes die at their current age.

This model was used to generate the plots presented in Figure 1, Figure 2D-E (using parameter estimates from Section 4.3) and Figures S4-S7.

### 4.3 Estimating the autocorrelation in the emergence rate of wild mosquito populations

To estimate autocorrelation in the emergence of wild mosquito populations from mosquito abundance data, we fitted an ordinary differential equation (ODE) based mosquito population model to EHT data. Age distribution data were not available for these mosquitoes. To identify the autocorrelation in the temporal trends in mosquito emergence, we fitted an ODE mosquito population model, with the same assumptions about mosquito survival and mortality, to the numbers of mosquitoes caught per experimental hut, *m*, irrespective of mortality and blood-feeding status. In EHTs, interventions and sleepers can correlate with differences in the total number of mosquitoes caught, but interventions and sleepers are rotated between experimental huts in a fully factorial design, so we do not explicitly account for these variables when estimating the temporal dynamics of mosquito emergence. We assume temporal changes in mosquitoes numbers, *M*, are determined by temporal changes in mosquito emergence only meaning,

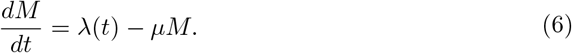

EHTs occur over relatively short timescales and mark-release-recapture studies indicate a lack of evidence for senescence in wild mosquitoes [45], so we assumed a constant per-capita mosquito mortality rate of *μ* = 0.132, in line with existing malaria transmission models [37]. Integrating equation with the assumption that the population-level emergence rate *λ*(*t*) is constant within each day gives,

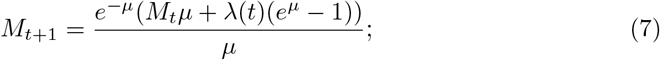

We fitted this model in a Bayesian framework assuming the daily mosquito counts per experimental hut are negative binomial distributed;

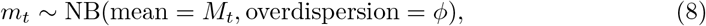

where *ϕ* is the overdispersion parameter, such that 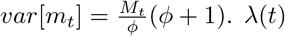 is the time-dependent mosquito emergence rate per day in the population as a whole and does not depend on the current mosquito population size; this quantity was fitted hierarchically using the lag-1 autoregressive model described in equation 5.

For each EHT, the posterior distributions of *M*_0_, *λ*_*t*_, *ϕ, h* and *σ* were estimated using the No U-turn Markov chain Monte Carlo (MCMC) sampler from Stan [57]. Wide normal priors were assumed: *M*_0_ ∼ N(15, 2.5), *ϕ*∼ N(1, 2.5), *h* ∼ N(0, 1) and *σ* ∼ N(0.5, 2.5). Convergence was assessed by visualising the trace plots and 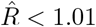 for all parameters. Results of the model fitting are presented in Figure 2A-C, with fitted parameter values shown in Figure S8.

We then simulated mosquito population dynamics using the IBM given the estimated constant daily emergence rates (*λ*(*t*)) and initial numbers of mosquitoes (*M*_0_) from the EHTs. To do so, we multiplied the fitted *λ*_*t*_ and *M*_0_ by the number of experimental huts to account for the mosquitoes in each experimental hut and inputted these values directly into the IBM. These results are presented in Figure 2D-E.

### 4.4 Compartmental mosquito model with ITNs

To model the effects of pyrethroid-only ITNs on the adult mosquito age distribution we used the *odin* and *odin*.*dust* R packages [58] to develop a discrete time stochastic compartmental approximation of an established differential equation larval and adult mosquito population model used in malaria transmission modelling [36–38]. Unlike the model outlined in equations 1-5 this model does not track individual mosquitoes but links the number of eggs and juvenile stages to the number of adult female mosquitoes. Within this model mosquitoes are classed as early stage larvae (*<u>E</u>*) late stage larvae (*L*), pupae (*P*) and female adult mosquitoes (*A*). We adapted this model, such that at each time step (*dt* = 1) the time-dependent numbers of births of new larvae is,

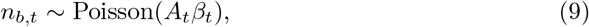

where *β*_*t*_ is the number of eggs laid per day per mosquito and *A*_*t*_ is the number of adult mosquitoes on day t (section S1.1). Male and female larval population dynamics are density dependent with the time-dependent changes in the mortality rates and carrying capacity, *K*(*t*),

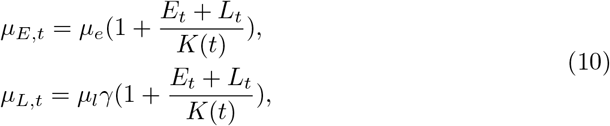

where *μ*_*e*_ = 0.0338 and *μ*_*l*_ = 0.0348 per-day are the per-capita daily mortality rates of early stage larvae and late stage larvae respectively. *μ*_*p*_ = 0.249 is the pupae mortality rate. Seasonality in the population dynamics occurs due to seasonal changes in the carrying capacity (section S1.1). Mosquitoes also transition due to development to other life stages; *d*_*el*_ = 6.64, *d*_*l*_ = 3.72 and *d*_*p*_ = 0.643 days are the development times of early stage larvae, late stage larvae and pupae respectively [36]. We assumed these rates are constant over a single time step and calculated the probability an event (death or development) for each life stage has occurs during that time step using an exponential CDF. The total rates of removal from each stage including both mortality and transition to other life stages were calculated as,

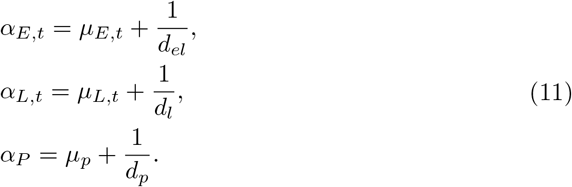

The numbers of mosquitoes that transition at each time step were then sampled thus,

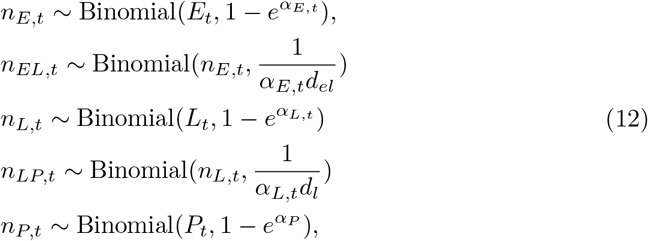

where *n*_*E,t*_, *n*_*L,t*_ and *n*_*P,t*_ are the numbers leaving the early larvae, late larve and pupae life stage compartments respectively; *n*_*EL,t*_ and *n*_*LP,t*_ are the numbers transitioning to the late larvae and pupae life stages respectively. Note that to calculate the numbers that develop to the next life stage and do not die we took a binomial sample from the total number transitioning with the probability determined by the proportion of the total rate account for by the development rate. The larval population dynamics are then calculated as,

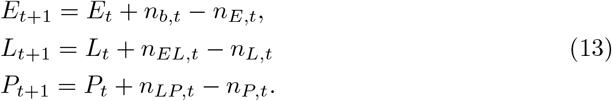

The adult female mosquito, *A*, compartments were stratified by age in time-steps; where *A*_*j,t*_, denotes the number of mosquitoes aged *j* (in dt) at time *t*. The ageing process was calculated thus,

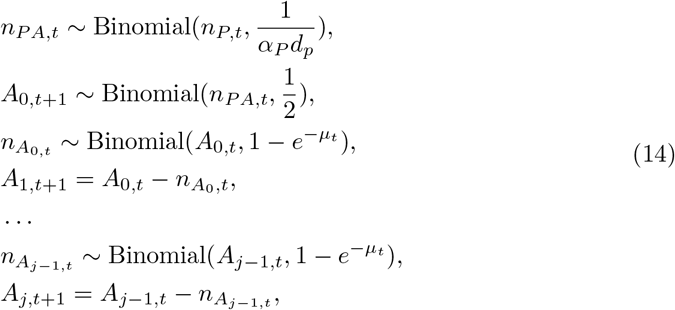

where *μ*_*t*_ is the per-capita adult mosquito mortality rate, which depends on the presence of ITNs (section S1.1). Note that the number of pupae developing into adults is drawn from a binomial sample with a 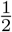 probability because only female adult mosquitoes are modelled.

In this analysis, we assume a population ITN usage of 0 or 100%, which does not change over time and an ITN entomological efficacy observed for pyrethroid-only ITNs with a mosquito population that is susceptible to pyrethroid insecticide (i.e. prior to the development of pyrethroid resistance) (supplementary information S1.1), assuming there is no insecticide resistance, 100% ITN coverage, and the proportions of anthropophagy (equal to 0.92), bites indoors (equal to 0.97) and bites when people are in bed (equal to 0.89) are high. This model is presented in Figure 3.

All code is available at https://github.com/IsaacStopard/mos_age_gono_IBM.

## 5 Acknowledgements

The work was supported by and the Bill and Melinda Gates Foundation (INV-078496), the Natural Environment Research Council (NERC; NE/P012345/1) and the UK Medical Research Council (MRC) Project Grant (MR/P01111X/1). IJS, ESS and TSC also acknowledge funding from the MRC Centre for Global Infectious Disease Analysis (reference MR/X020258/1), funded by the United Kingdom MRC. This United Kingdom funded award is carried out in the frame of the Global Health EDCTP3 Joint Undertaking. The funders had no role in study design, data collection and analysis, decision to publish or preparation of the manuscript. The authors would like to thank Nakul Chitnis and Maria-Gloria Basáñez for comments on early versions of the work.

